# Antibiograms and Molecular Characterization of Drug Resistance of *Mycobacterium abscessus* complex from Patients with Multidrug-Resistant Pulmonary Tuberculosis (MDR TB) Infection

**DOI:** 10.1101/864447

**Authors:** Kenneth A. Bongulto, Concepcion F. Ang, Esperanza C. Cabrera

## Abstract

In the Philippines, acid fast bacilli positive sputum samples commonly treated as TB due to *Mycobacterium tuberculosis* (MTB) complex. However, *Mycobacterium abscessus* (MAB) complex is often found in MTB cultures, or in patients confirmed negative for TB through sputum microscopy and culture. Hence, patients with MAB infections are mistakenly prescribed six-month anti-TB treatments. In this study, MAB complex isolates from MDRTB patients were identified and further sub-speciated using the *mass* 3210 gene. Antimicrobial susceptibility was tested using broth microdilution and resistance genes *erm*(41), *rrs, rrl, gyr*A, and *gyr*B were studied for mutations. Majority were susceptible to amikacin, azithromycin, clarithromycin, and moxifloxacin [MAB: 100%, 100%, 100%, 81.8%, respectively; *M. massiliense* (MAM): 100%, 100%, 100%, 60%, respectively]. 50% MAM and 63.6% MAB were susceptible to cefoxitin; 60% MAM and 45.5% MAB were susceptible to ciprofloxacin; 72.7% MAB, and 10%MAM were susceptible to doxycycline. Inducible resistance to azithromycin and clarithromycin was found in 27.3%MAB and 30% MAM. 42.9% MAB complex isolates were MDR. Macrolide resistant MAB and MAM had T28 sequevar, showing functional *erm*(41) responsible for inducible resistance. Unexpectedly, full length *erm*(41) was found in MAM. The*rrl* gene in these isolates showed no point mutations, indicating T28 sequevar as cause of inducible resistance. All fluoroquinolone resistant isolates showed Ala-83 in *gyr*A fluoroquinolone resistant-dependent region (QRDR) and Arg-447 and Asn-464 in *gyr*B QRDR. These are associated with resistance to the drug.

## INTRODUCTION

In the Philippines, one of the most common rapidly growing mycobacteria (RGM) from patients with multiple drug resistant tuberculosis (MDR-TB) infection is *M. abscessus* complex. This non-tuberculous *Mycobacterium* (NTM) species is usually found in TB culture, either growing alongside with *M. tuberculosis* or after the patient has been confirmed negative for pulmonary tuberculosis through direct sputum smear microscopy (DSSM) and TB culture. Pulmonary diseases caused by the *M. abscessus* complex are extremely hard to manage for these species are found to be resistant to anti-tuberculous agents. Patients with *M. abscessus* infection do not receive appropriate treatment for they are considered to have chronic TB and MDR-TB. As a result, patients are prescribed a six-month anti-TB treatment which is not appropriate for *M. abscessus* infections. Interestingly, there is no known study in the Philippines regarding the antimicrobial resistance profile of *M. abscessus* complex. NTM studies are overshadowed by pulmonary TB research studies and therefore, clinicians are not guided on the treatment for NTM infections. Moreover, detection of NTM species is not part of the routine procedure in the clinical microbiology laboratory. Treatment failure to both first-line anti-TB drugs rifampicin and isoniazid is linked to infection with multidrug-resistant *M. tuberculosi*s (MDR-TB). Gler *et al.* (2012) showed very high rates (83% to 97%) of MDR-TB infections in the Philippines in 2012. Because MDR TB is often associated with the occurrence of *M. abscessus* complex in clinical samples, the present study aimed to characterize local isolates of *M. abscessus* complex from patients with multidrug-resistant pulmonary tuberculosis in terms of their antimicrobial susceptibility profiles and determine the genetics of the antibiotic resistance. More specifically, this study aimed to determine the phenotypic characteristics of *M. abscessus* complex using colony morphology, growth rate, and pigmentation; to determine subspecies of the *M. abscessus* complex isolates using amplification and sequencing of the *mass_3210* gene; to assess susceptibility of *M. abscessus* complex to antimicrobials (clarithromycin CLR, azithromycin AZM, amikacin AMK, ciprofloxacin CIP, moxifloxacin MXF, cefoxitin FOX, and doxycycline DOX) commonly used against it; and to determine the mechanism of antibiotic resistance by conducting sequence analysis of the drug resistance related genes [*erm*(41), *rrl, rrs, gyr*A, and *gyr*B] in selected isolates.

## MATERIALS AND METHODS

### Study Site and Bioethical Clearance

All procedures involving handling of infectious materials (culture, DNA extraction, NTM identification, and drug susceptibility tests) were conducted in the P3 laboratory of the National Center for Pulmonary Research, Lung Center of the Philippines. The study sought approval form the Institutional Ethics Review Board of the Lung Center of the Philippines. The requirement for informed consent was waived since the isolates from the stock cultures to be studied were anonymized, but bioethical clearances from the De La Salle University – Manila, and the Lung Center of the Philippines were procured.

### Study Isolates and Identification of *Mycobacterium abscessus*

The sample size computation for an interval estimate of population was used to determine the required sample size of *M. abscessus* complex isolates for the study. This is as follows: n= t^2^ × p (1-p) / m^2^, where *n* is the required sample size, t is the confidence level at 95% (standard value of 1.96), p is the estimated prevalence of NTM, and m is the relative precision (Charan & Biswas, 2013). The prevalence rate used for the sample size computation was from the report of Siapno *et al*. (2016) on a retrospective study of 6,886 specimens from NTM infection in a tertiary hospital in the Philippines, which was 2.28%. The relative precision used was one-fifth of the prevalence which was 0.00456. Since the prevalence rate was small, the precision of 5% seemed to be inappropriate. A conservative choice would be one-fourth or one-fifth of prevalence as the amount of precision in the case of small prevalence (Pourhoseingholi *et al*., 2013). The computed sample size of 20 was based on NTM, and the study adapted this as 20 *M. abscessus* complex. One hundred seventeen (117) NTM clinical isolates collected from patients with multiple drug-resistant TB infections and were confirmed MPT64 antigen negative were obtained from the Programmatic Management of Drug Resistant Tuberculosis (PMDT) TB culture located at the National Center for Pulmonary Research, Lung Center of the Philippines from which these 20 isolates identified to be *M. abscessus* complex were taken.

### Phenotypic characterization of *Mycobacterium abscessus* complex

Freshly grown 5 to 7-day old cultures of *M. abscessus* complex were subjected to phenotypic characterization. Colony morphology was determined by observing the colonies grown on Ogawa medium. There are two possible morphotypes of *M. abscessus* complex, namely: the smooth and rough colonies. Growth rate was also determined by checking the presence of visible colonies on Ogawa media on a daily basis. Pigmentation was determined by observing the color of the colonies on the Ogawa medium. *Mycobacterium abscessus* complex is comprised of non-photochromogens. All isolates were tested using the Ziehl Neelsen method to confirm the presence of acid fast bacilli (AFB) and to check the purity of the culture. More so, Kudoh method was conducted in cultures with contaminants. This involved adding 5% NaOH to the culture of *M. abscessus* complex grown in 7H9 broth in a ratio of 1:1 (v/v) The processed isolates were re-incubated at 37°C for 5 to 7 days. AFB smear was conducted again, to check for purity and to confirm presence of acid-fast bacilli.

### Identification of subspecies of *Mycobacterium abscessus* complex

One hundred seventeen (117) NTM clinical isolates were screened for the identification of *M. abscessus* and *M. massiliense* isolates. All isolates were grown in Middlebrook 7H9 broth enriched with albumin dextrose catalase (ADC). These were incubated at 37°C for 5 to 7 days or until turbidity was observed and checked for purity. New stock cultures were prepared in Ogawa butt-slants to serve as working stocks, and in 10% glycerol at −70°C.

For identification, DNA extraction was done on freshly grown 5 to 7 day-old cultures using Bio-Rad Chelex resin (Al-Mutairi *et al*., 2011) as follows: A loopful of mycobacterial isolate was suspended in a 200 μL solution containing 40mg Chelex100 and 100 mL of water. The resulting material was kept at 95°C for 20 minutes. It was centrifuged for 15 minutes at 12,000 x g. The supernatant was transferred to a sterile Eppendorf® tube and was used as the source of DNA for the one-step multiplex PCR assay. The one-step multiplex PCR assay designed by Chae *et al.* (2017) was used (1) to discriminate pan-mycobacterial from non-mycobacterial species by amplifying the16S rRNA gene, (2) to distinguish between MTB complex and NTM species in mycobacteria by amplifying the *rv0577*, (3) to identify *M. tuberculosis* by amplifying the RD9, (4) to identify *M. tuberculosis* Beijing family by amplifying the *mtbk_20680*, and (5) to identify the five major NTM species and subspecies by amplifying IS*1311* (*M. avium*), DT1 (*M. intracellulare*), *mass_3210* (*M. abscessus* and *M. massiliense*), *mkan_rs12360* (*M. kansasii*).

The PCR mixture was comprised of 1) 2x Prime TAq Premix (Genet Bio., Ltd. Daejeon, South Korea) containing Prime *Taq* DNA polymerase 1 unit/10ul, 2X reaction buffer, 4mM MgCl_2_, enzyme stabilizer, sediment, loading dye (pH 9.0), and 0.5 mM of each dATP, dCTP, dGTP, and dTTP; 2) 10 pmole of each of the following forward and reverse primers of *rv0577*F (5’-GAG ATA CTC GAG TGG CGA A-3’), *rv0577*R (5’-CAA CGC GAC AAA CCA CCT AC-3’), *DT*1F (5’-AAG GTG AGC CCA GCT TTG AAC TCC A-3’), *DT*1R (5’-GCG CTT CAT TCG CGA TCA TCA GGT G-3’), mtbk*_20680*F (5’-TTA TGC CAG AAA TAC ACC CGC G-3’), mtbk*_20680*R (5’-AAT CGC GGG CTT GTG GCT AC-3’), 16S rRNAF (5’-GAG ATA CTC GAG TGG CGA AC-3’), 16S rRNAR (5’-CAA CGC GAC AAA CCA CCT AC-3’), RD9F (5’-GTG TAG GTC AGC CCC ATC C-3’), RD9R (5’-GTA AGC GCG TGG TGT GGA-3’), IS*1311*F (5’-TCG ATC AGT GCT TGT TCG CG-3’), IS*1311*R (5’-CGA TGG TGT CGA GTT GCT CT-3’), *mass_3210*F (5’-GCT TGT TCC CGG TGC CAC AC-3’), *mass_3210*R (5’-GGA GCG CGA TGC GTC AGG AC-3’), *mkan_rs12360*F (5’-ACA AAC GGT GTG TCG CAA TGT GCC A-3’), and *mkan_rs12360*R (5’-TGT CGA GCA GAC GTT CCA GGA CGG T-3’), respectively; 3) 2 μl DNA template; and 4) sterile deionized distilled water. The cycling condition included an initial denaturation step at 95°C for 10 minutes, then 30 cycles at 96°C for 45 seconds, 61.5°C for 45 seconds, and 72°C for 40 seconds, followed by 72°C for 10 minutes for the final extension. The amplicons were analyzed using 2% gel for 40 minutes at 100 V in 0.5X TBE buffer. The gels were visualized using the Bio-Rad ImageLab gel reader (Bio-Rad Laboratories Inc. USA).

### Antimicrobial Susceptibility Testing

The mycobacterial isolates were cultured on Ogawa medium at 37°C for 5 to 7 days. Suspensions were prepared by aseptically sweeping the confluent portion of growth on the Ogawa medium with a sterile loop. Growth on the sterile loop was transferred to 4.5 mL of sterile water containing glass beads (7 to 10 3-mm beads) until the turbidity matched the 0.5 McFarland standard with an approximate organism density of 1×10^5^ colony forming units ml^-1^ (CFU/mL). The final inoculum with 5×10^2^ CFU/mL was prepared by transferring 50 *μ*l of the suspension to a tube containing 10 ml of cation adjusted Mueller-Hinton broth (CLSI M24-A2, 2011).

Antimicrobial susceptibility testing of *M. abscessus* complex isolates was performed using the gold standard assay recommended by the Clinical Laboratory Standards Institutes (CLSI M24-A2, 2011) which is the microdilution technique. BBL cation adjusted Mueller-Hinton (CAMH) Broth (BD, Franklin Lakes, NJ, USA) with 5% OADC was used as the medium for the tests. Tests on the mycobacterial strains were conducted in 96-well microplates. The broth microdilution method is traditionally set up as two-fold dilutions. The following working ranges were used for the mycobacteria: the concentration range for amikacin, cefoxitin, and azithromycin was 0.25 to 256 *μ*g/mL; the concentration range for clarithromycin, ciprofloxacin, doxycycline, and moxifloxacin was 0.0625 to 64 *μ*g/mL (CLSI M24-A2, 2011). All antimicrobial drugs used for the antimicrobial susceptibility test were procured from Sigma Aldrich (Merck Group, USA). To determine the amount of antimicrobial agent powder needed for a standard solution, the formula outlined in the CLSI M24-A2, 2011 was used.

The thoroughly mixed antimicrobial dilutions (100 *μ*l) were dispensed into each well, except for the 11^th^ well, which is a drug-free well for bacterial growth control. The 1^st^ well contained 200 *μ*l of CAMHB and served as the negative control. A total of 100 *μ*l of the final inoculum was dispensed in the second well up to the 11^th^ well. The final volume in each well was 200 *μ*l. Reference strain of antimicrobial susceptible *M. peregrinum* ATCC^®^ 700686 was tested along with the test clinical isolates to serve as the quality control. The inoculated 96-well microplate were sealed and incubated at 37°C. The MIC values for amikacin, cefoxitin, doxycycline, ciprofloxacin, moxifloxacin, azithromycin and clarithromycin were read on the third day after the inoculum was added. The MICs of azithromycin and clarithromycin for isolates showing susceptible results on the third day were read again on the 7^th^ day and 14^th^ day to determine the presence of inducible resistance to the macrolides (Nie *et al.*, 2014). Inducible resistance to macrolides was concluded once the mycobacterial strains showed azithromycin or clarithromycin susceptibility on the third day and resistance on the 7th or 14^th^ day (Lee *et al*., 2014). Thirty microliters of a freshly prepared 0.01% of 10X resazurin (Accumed International, Westlake, Ohio) reagent were added to all wells in the microplate. A blue color in the well was interpreted as no growth, while a pink color was scored as growth. The MIC was defined as the lowest drug concentration which prevented the color change from blue to pink (Franzblau *et al.*, 1998). Each test was done in triplicate.

### PCR amplification of *rrl, erm*(41), *rrs, gyrA*, and *gyrB* genes

Sequencing of *rrl* and *erm*(41) genes was conducted to compare and analyze the characteristics of the susceptible and resistant *M. abscessus* strains to AZM and CLR, respectively. The primer sets are as follows: *erm*(41)F (5’-GAG CGC CGT CAC AAG ATG CAC A-3’), *erm*(41)R (5’-GAC TTC CCC GCA CCG ATT CCA C-3’), *rrl*F (5’-GTA GCG AAA TTC CTT TGT CGG-3’), and *rrl*R (5’-TTC CCG CTT AGA TGC TTT CAG-3’) (Nash *et al*., 1995; Nash *et al*., 2009). The primer sets for the amplification of the amikacin resistance-related gene are *rrs*F (5’-CAG TAC AGA GGG CTG CGA ACG-3’) and *rrs*R (5’-AAG GAG GTG ATC CAG CCG CA-3’) (Prammananan *et al.*, 1998). To characterize resistant *M. abscessus* strains to CIP and MXF, sequence analysis of the fluoroquinolone resistant-dependent region (QRDR) of the *gy*r*A* and *gyrB* genes was conducted. The primer sets are: *gyr*AF (5’-GGG CAT CTA AAG CCG CTG AGA-3’), *gyr*AR (5’-GAC GAT GGC GCG CTG ACG T-3’), *gyr*BF (5’-GCA GAT GCT AAA ACG GTT GTG A-3’) and *gyr*BR (5’-CTC GTA AGT ACG ACG GCA CAA-3’) (Guillemin *et al.*, 1998).

## RESULTS and DISCUSSION

### Phenotypic Characterization of *Mycobacterium abscessus* complex

The *M. abscessus* complex isolates were characterized based on colony morphology, growth rate, and pigmentation. The isolates tended to aggregate and form biofilms on the third day of incubation in 7H9 broth medium incubated at 37°C. Biofilms were visible both on the surface and at the bottom of the tubes. The enriched cultures of *M. abscessus* complex from 7H9 medium were grown in Ogawa butt slants to determine the proportion of smooth and rough colonies. Growth on the surface of the Ogawa medium was observed on the third day of incubation at 37°C.

The phenotypic characteristics of the 21 *M. abscessus* complex isolates are shown in Figures 1 and 2, and Table 1. Smooth colonies of *M. abscessus* and *M. massiliense* were moist, shiny, and round, while the rough colonies were dry, waxier, and wrinkled. The smooth colony of *M. abscessus* complex expresses glycopeptidolipid (GPL) on its cell wall and forms biofilms, while rough colony of *M. abscessus* complex does not form biofilms due to minimal amounts of GPL (Howard *et al*., 2006; Byrd & Lyons, 1999). Moreover, rough colony of *M. abscessus* is associated with virulence compared to the smooth colony of *M. abscessus*. In the present study, a total of 7 isolates (63.6%) of *M. abscessus* and 3 (30.0%) *M. massiliense* isolates were characterized to have smooth colonies, while 4 (36.4%) *M. abscessus* and 7 (70.0%) *M. massiliense* isolates produced rough colonies. All *M. abscessus* and *M. massiliense* isolates were buff – colored and were categorized as nonphotochromogens, i.e., they were non-pigmented whether they were grown in the dark or in the presence of light. All (100%, 21/21) isolates that grew in Ogawa medium were acid fast. NTM appears as shorter rods compared to *M. tuberculosis* when viewed microscopically.

**Table 1.** Phenotypic characteristics of *M. abscessus* and *M. massiliense* isolates from patients with MDR TB infections.

| Study ID | AFB Smear | Growth Rate | Pigmentation | Colony Morphology |
| --- | --- | --- | --- | --- |
| MAB01 | + | Day 3 | buff | smooth |
| MAB02 | + | Day 3 | buff | rough |
| MAB03 | + | Day 3 | buff | rough |
| MAB04 | + | Day 3 | buff | smooth |
| MAB05 | + | Day 3 | buff | smooth |
| MAB06 | + | Day 3 | buff | smooth |
| MAB07 | + | Day 3 | buff | rough |
| MAB08 | + | Day 3 | buff | rough |
| MAB09 | + | Day 3 | buff | smooth |
| MAB10 | + | Day 3 | buff | smooth |
| MAB11 | + | Day 3 | buff | smooth |
| MAS01 | + | Day 3 | buff | rough |
| MAS02 | + | Day 3 | buff | rough |
| MAS03 | + | Day 3 | buff | rough |
| MAS04 | + | Day 3 | buff | rough |
| MAS05 | + | Day 3 | buff | smooth |
| MAS06 | + | Day 3 | buff | smooth |
| MAS07 | + | Day 3 | buff | rough |
| MAS08 | + | Day 3 | buff | smooth |
| MAS09 | + | Day 3 | buff | rough |
| MAS10 | + | Day 3 | buff | rough |

MAB = *M. abscessus*; MAS = *M. massiliense*

**Figure 1.**
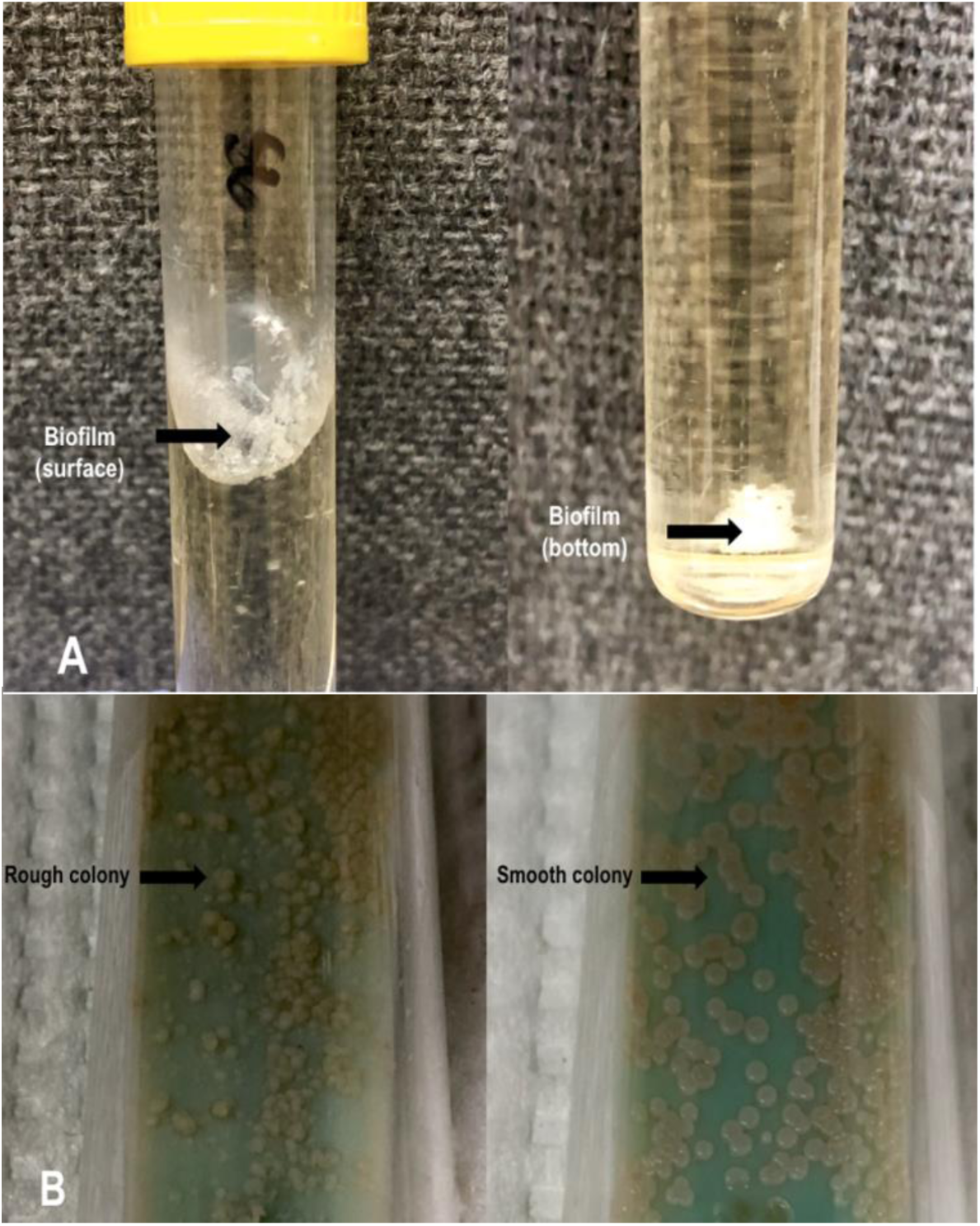
Biofilm formation of *M. abscessus* complex in 7H9 medium (A). Colony morphology of *M. abscessus* complex isolates showing two morphotypes: rough and smooth colonies (B). All isolates are nonphotochromogens (non-pigmented).

**Figure 2.**
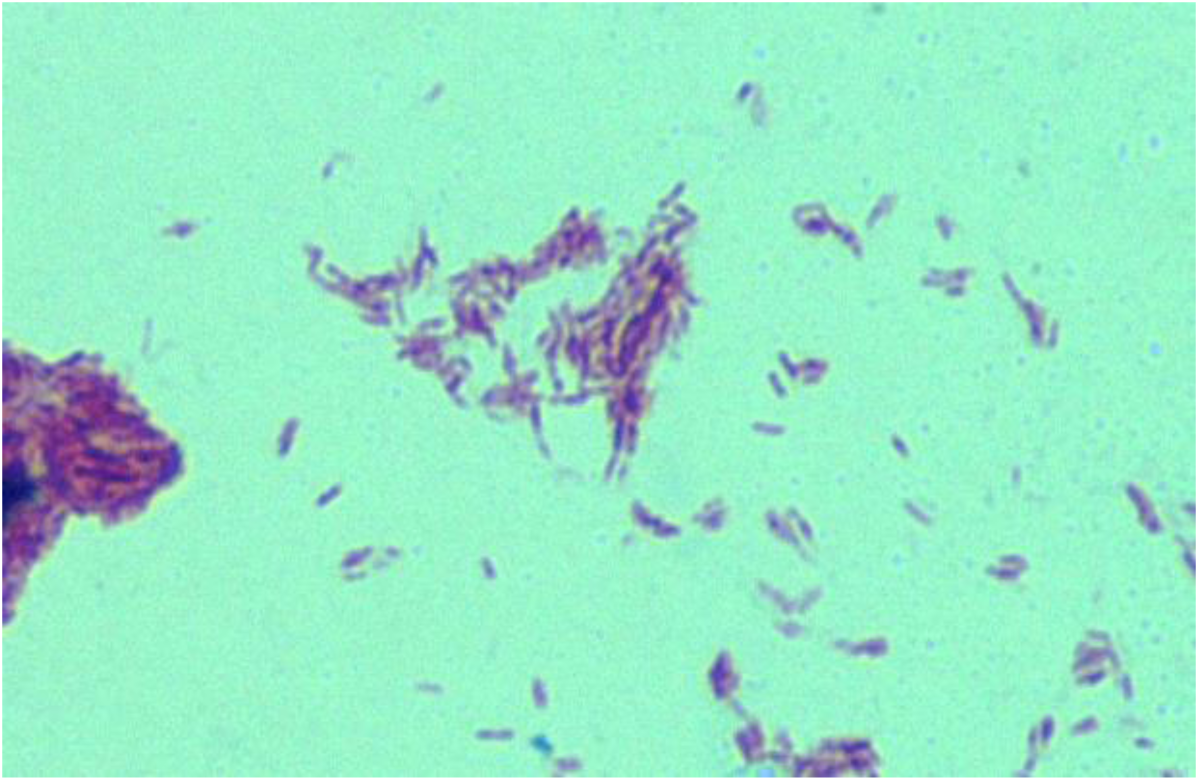
Positive AFB smear result under 1000x magnification, with immersion oil.

### Subspecies identification of *Mycobacterium abscessus* complex isolates

The subspecies of the 117 clinical isolates that were found to be negative for MPT64 were identified using multiplex PCR. Among the 117 clinical isolates, 11 (9.4%) were identified as *M. abscessus* and 10 (8.5%) were *M. massiliense* (Figure 4). The remaining isolates were identified as *M. tuberculosis* (15%, 17/114), other NTM species (53.8%, 63/117), and other microorganisms (14%, 16/114). It is also noteworthy that all 117 isolates were found negative using the MPT64 antigen kit, but some were identified as *M. tuberculosis* using the one-step multiplex PCR assay. These results may infer that mixed colonies of *M. tuberculosis* and NTM might be present in the stock cultures from the bank specimens. Purification was conducted on all identified *M. abscessus* complex isolates. Positive controls of *M. abscessus* and *M. massiliense*, which were clinical isolates from the bank specimens in the National Center for Pulmonary Research – TB laboratory that were previously identified by reverse blot hybridization assay were tested along with the *M. abscesus* complex test isolates.

**Figure 3.**
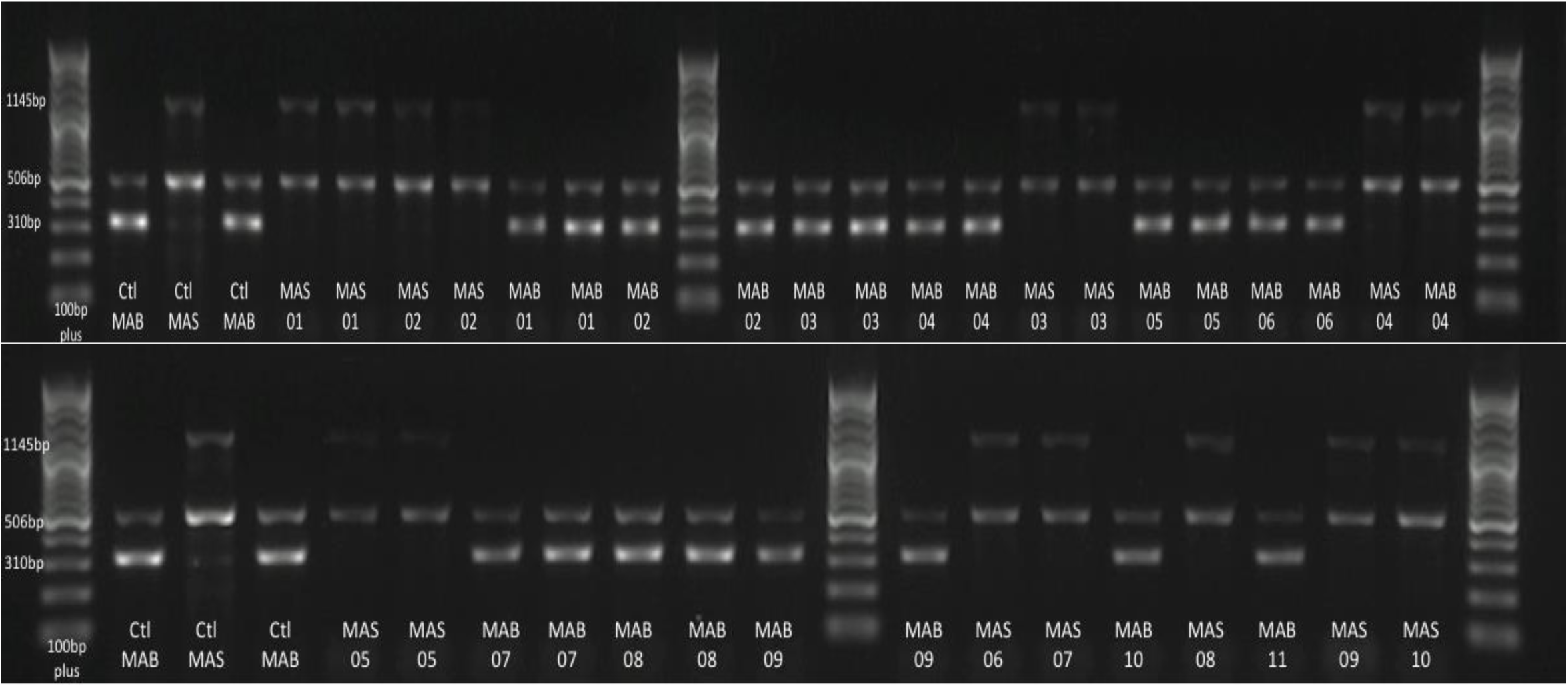
One-step Multiplex PCR Assay results using primers for *mass_*3210 gene. Molecular marker (100 plus bp); Ctrl MAB: *M. abscessus* control (310bp *mass_*3210 amplicon) and the *Mycobacterium*-specific16S rRNA internal control (506bp); Ctrl MAS: *M. massiliense* control (1145bp *mass_*3210 amplicon) and the *Mycobacterium*-specific 16S rRNA internal control (506bp). MAS = *M. massilience;* MAB = *M. abscessus.*

**Figure 4.**
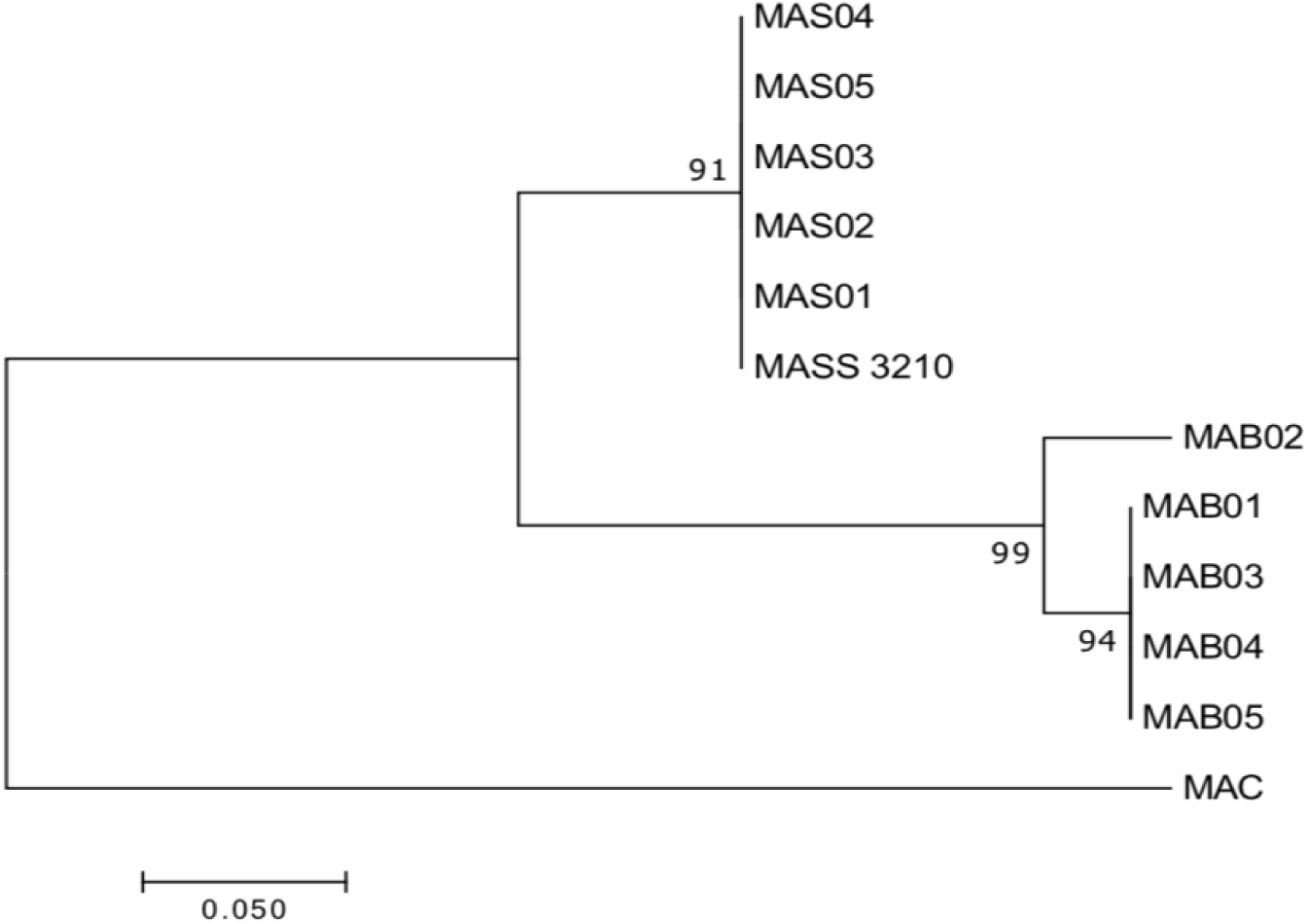
Neighbor-joining tree based on *mass*_3210 sequences. Dendrogram showing the clustering of *M. abscessus* isolates (MAB01-MAB05) and *M. massiliense* isolates (MAS01-MAS05) together with *mass*_3210. The *IS*_1311 sequence which is specific to *M. avium* complex (MAC) was used as the outgroup.

All mycobacteria showed the 506bp band corresponding to the 16S rRNA internal control which was specific only to all mycobacterial species. *M. abscessus* isolates showed the expected 310bp band and *M. massiliense* showed the 1145bp band using the primers for *mass*_3210. The *mass_*3210 is a specific gene previously identified and used to discriminate between *M. abscessus* complex species (Chae *et al.*, 2017). Further analysis of the *mass_*3210 aligned sequences of representative isolates showed two clustered groups: MAB01-MAB05 and MAS01-MAS05 (Figure 4). This result implies that MAB01-MAB05 isolates clustered together, showing that they belonged to one subspecies (*M. abscessus*). On the other hand, MAS01-MAS05 isolates grouped together, showing that they belonged to *M. massiliense* subspecies.

### Drug-resistance Profile of *Mycobacterium abscessus* complex

Results of the antimicrobial susceptibility assay of the *M. abscessus* complex are shown in Table 2. The present study showed that amikacin was the most active agent against all *M. abscessus* and *M. massiliense* isolates with an MIC of <1μg/ml for both groups. These are in conformity with the results of the study of Nie *et al.* in 2014 which showed that the susceptibility rate of *M. abscesuss* to AMK was 98%. Likewise, Park *et al.* (2008) reported a 99% susceptibility rate of *M. abscessus* to AMK, while Kim *et al.* (2015) reported the susceptibility rate of *M. abscessus* and *M. massiliense* to AMK as 91.2% and 100%, respectively. The susceptibility of all isolates to amikacin in the present study may be attributed to the current treatment regimen for MDR-TB in the country which is comprised of kanamycin and not amikacin. Amikacin resistant isolates are not positively selected for to survive and thus are not disseminated.

**Table 2.**
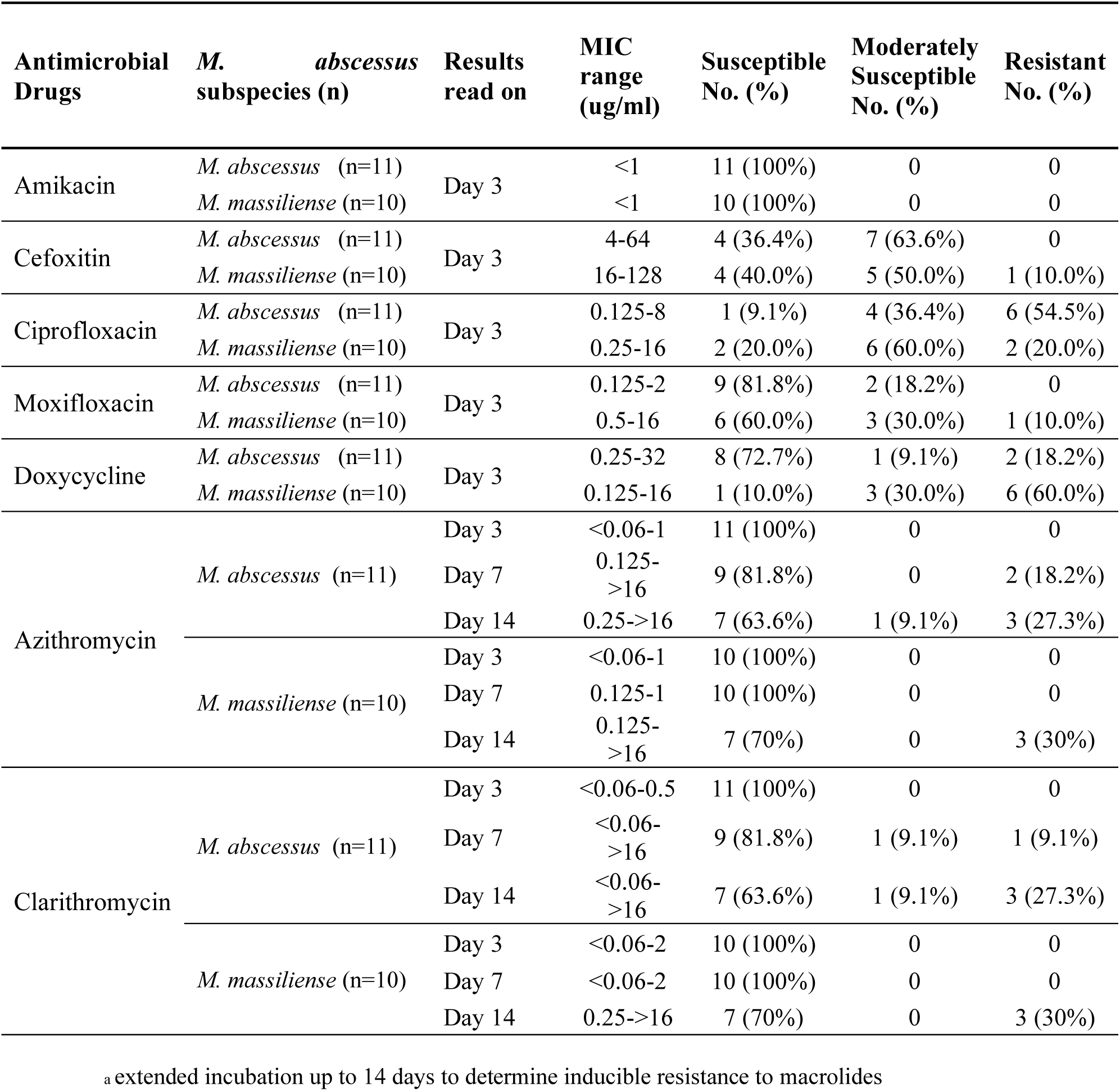
Antimicrobial susceptibility test results of *M. abscessus* and *M. massiliense* using broth microdilution method against seven (7) antimicrobial drugs.

On the other hand, FOX was shown to be less potent *in vitro* against *M. abscessus* and *M. massiliense* isolates compared to amikacin. Although there were no *M. abscessus* isolates that were resistant to the drug, and a low 10% of the *M. massiliense* were resistant to the agent, 63.6% of the *M. abscessus* and 50% of the *M. massiliense* isolates were only moderately susceptible to cefoxitin. The remaining 36.4% and 40% of the *M. abscessus* and *M. massiliense* isolates were susceptible to it. The results are comparable to those of Nie *et al.* (2014) that showed 0% resistant *M. abscessus* isolates, while 53% were moderately susceptible. Koh *et al.* (2010) likewise showed no resistant isolates of *M. abscessus* and only 1% resistance rate in *M. massiliense* against FOX. Lee *et al.* (2015) reported resistance rate of 1.3% in *M. abscessus* subsp. *massiliense* and 0% for *M. abscessus* subsp. *abscessus* against FOX.

Susceptibility rates of *M. abscessus* complex isolates in the present study showed that MXF has better *in vitro* activity against *M. abscessus* (81.8% susceptible) and *M. massiliense* (60% susceptible) compared to CIP against *M. abscessus* (9.1% susceptible) and *M. massiliense* (20% susceptible). This is consistent with other studies showing that MXF has better *in vitro* activity against *M. abscessus* and *M. massiliense* compared to CIP (Kim *et al.*, 2015; Lee *et al.*, 2014; Koh *et al.*, 2010). On the other hand, DOX was found to be more potent against *M. abscessus* (72.7% susceptible) isolates compared to its activity against *M. massiliense* (10% susceptible). The resistance rate of *M. massiliense* (60%) in the present study is compatible with the findings of Koh *et al.* (73%) and Lee *et al.* (73.4%).

Macrolides, specifically, AZM and CLR are the drugs of choice for the treatment of infections caused by the *M. abscessus* complex. The present study reported 100% susceptibility rate for both *M. abscessus* and *M. massiliense* isolates on the 3^rd^ day of incubation. However, inducible resistance to both CLR and AZM were found in *M. abscessus* MAB04 with MICs of >16ug/ml for both drugs on the 7^th^ day of incubation, while *M. abscessus* MAB07 showed inducible resistance to AZM with an MIC of 16μg/ml on the 7^th^ day, and inducible resistance to CLR on the 14 th day (MIC 16μg/ml). *Mycobacterium abscessus* isolate MAB09 showed inducible resistance to both CLR (>16μg/ml) and AZM (>16μg/ml) on the 14^th^ day of incubation. Inducible resistance in *M. abscessus* was expected since they have a functional erm(41) gene (Kim *et al.*, 2015; Koh *et al.*, 2010; Nash *et al.*, 2009). On the other hand, three *M. massiliense* isolates (MAS04: >16μg/ml, MAS06: >16μg/ml, MAS10: >16μg/ml) exhibited inducible resistance to CLR and AZM on the 14^th^ day. This is contrast with previous studies (Kim *et al.*, 2015; Koh *et al.*, 2010) which demonstrated that inducible resistance to CLR was only present in *M. abscessus* and not in *M. massiliense.* Nevertheless, our results support the findings of Li *et al*. (2017) and Shallom *et al*. (2013) showing that *M. massiliense* exhibited an inducible resistance to CLR. Lastly, the MIC of the test drugs for the susceptible reference *M. peregrinum* ATCC 700686 used for quality control were all in accordance with the tentative quality control ranges for rapidly growing mycobacteria as indicated in the CLSI M24-A2 (2011) guidelines.

The resistance phenotypes of *M. abscessus* complex isolates are summarized in Table 3. Overall, 42.9% (9/21) of the *M. abscessus* complex isolates were found MDR; 14.3% of which were *M. abscessus* isolates and 28.6% of were *M. massiliense* isolates. MDR is defined as resistance to at least one agent in three or more antimicrobial categories (Magiorakos *et al.*, 2012).

**Table 3.** Resistance phenotypes of *M. abscessus* and *M. massiliense* in the study.

| Study ID | Subspecies Identification | Antimicrobial Agents |  |  |  |  |  |  | MDR* |
| --- | --- | --- | --- | --- | --- | --- | --- | --- | --- |
|  |  | AZM | CLR | AMK | FOX | CIP | MXF | DOX |  |
| MAB01 | *M. abscessus* | S | S | S | M | R | S | M | + |
| MAB02 | *M. abscessus* | S | S | S | M | M | S | S | - |
| MAB03 | *M. abscessus* | S | S | S | M | S | S | S | - |
| MAB04 | *M. abscessus* | IR | IR | S | M | S | S | S | - |
| MAB05 | *M. abscessus* | S | S | S | M | R | S | S | - |
| MAB06 | *M. abscessus* | S | S | S | M | R | S | S | - |
| MAB07 | *M. abscessus* | IR | IR | S | S | M | S | S | - |
| MAB08 | *M. abscessus* | S | S | S | S | R | S | S | - |
| MAB09 | *M. abscessus* | IR | IR | S | M | R | M | R | + |
| MAB10 | *M. abscessus* | S | S | S | M | R | S | S | - |
| MAB11 | *M. abscessus* | S | S | S | M | M | M | R | + |
| MAS01 | *M. massiliense* | S | S | S | S | M | S | M | - |
| MAS02 | *M. massiliense* | S | S | S | M | S | S | M | - |
| MAS03 | *M. massiliense* | S | S | S | S | S | M | R | - |
| MAS04 | *M. massiliense* | IR | IR | S | M | S | S | M | + |
| MAS05 | *M. massiliense* | S | S | S | M | S | M | R | + |
| MAS06 | *M. massiliense* | IR | IR | S | R | R | M | R | + |
| MAS07 | *M. massiliense* | S | S | S | M | M | M | S | - |
| MAS08 | *M. massiliense* | S | S | S | M | R | R | R | + |
| MAS09 | *M. massiliense* | S | S | S | M | M | S | R | + |
| MAS10 | *M. massiliense* | IR | IR | S | M | M | S | R | + |

Antimicrobial agents tested AZM (Azithromycin), CLR (Clarithromycin), AMK (Amikacin), FOX (Cefoxitin), CIP (Ciprofloxacin), MXF (Moxifloxacin), DOX (Doxycycline). The letter S is for susceptible, M indicates moderate susceptible, R is for resistant, and IR stands for inducible resistant isolate. The hyphen indicates that the isolate is not MDR while the (+) sign indicates that the isolate is MDR. Multidrug-resistant (MDR) is defined as resistance to at least one agent in three or more antimicrobial categories (Magiorakos *et al.*, 2012).

Although the MDR percentage obtained in the study is lower than the 97.2% MDR report of Candido *et al.* (2104), where *M. abscessus* complex isolates were found to be resistant to five or more antimicrobial agents, the emergence of multidrug-resistant *M. abscessus* complex isolates in the country is of serious concern which impacts on the current treatment regimen given to patients with *M. abscessus* infection. This calls for the need to conduct antimicrobial susceptibility testing of clinical isolates of *M. abscessus* complex for better management of the patients, and for the prevention of the selection for resistant strains to survive and be disseminated.

### Molecular Characterization of Drug-resistance of *Mycobacterium abscessus* complex

In the present study, the *erm*(41) gene of all the *M. abscessus* and *M. massiliense* isolates with inducible resistance was amplified. All three *M. abscessus* isolates (MAB04, MAB07, MAB09) showing inducible resistance to CLR and AZM were found to have a full-length *erm*(41) gene of 892bp. These isolates with the full-length *erm*(41) gene have a T28 sequevar which is associated with inducible resistance to macrolides. Such resistance can affect the current treatment regimen of a patient, thus contributing to the hurdles towards successful treatment. Interestingly, all three *M. massiliense* isolates (MAS04, MAS06, MAS10) with an MIC of >16ug/ml harbored a full-length *erm*(41) gene that is present only in *M. abscessus* (Nash *et al.*, 2009), instead of the expected truncated *erm*(41) gene. This finding supports the results of other studies (Lipworth *et al.*, 2018; Shallom *et.*, 2013), which also reported the presence of a functional *erm*(41) gene in *M. massiliense* leading to a resistant phenotype. The mutation patterns of the macrolide-resistant *M. abscessus* complex isolates are summarized in Table 4. Caution should thus be practiced in using the *erm*(41) gene in identifying subspecies of *M. abscessus* complex.

**Table 4.** Single nucleotide polymorphisms (SNPs) in the *erm*(41) gene associated with macrolide-resistant isolates of *M. abscessus* and *M. massiliense*

| Study ID | T28<br><u>TGG</u><br>(Trp) | T159<br><u>GGT</u><br>(Gly) | G168<br><u>GTG</u><br>(Val) | T231<br><u>GAT</u><br>(Asp) | A238<br><u>ATA</u><br>(Ile) | G249<br><u>GCG</u><br>(Ala) | C253<br><u>CTA</u><br>(Leu) | A255<br><u>CTA</u><br>(Leu) | A312<br><u>CAA</u><br>(Glu) | A330<br><u>ATA</u><br>(Ile) | T336<br><u>AGT</u><br>(Ser) | A414<br><u>CGA</u><br>(Arg) |
| --- | --- | --- | --- | --- | --- | --- | --- | --- | --- | --- | --- | --- |
| MAB4 | - | C | - | C | G<br>(Val) | A | A (Ile) | T (Ile) | C<br>(His) | C | C | G |
| MAS4 | - | - | C | - | - | - | - | G | - | - | - | - |
| MAB7 | - | - | - | - | - | - | - | G | - | - | - | - |
| MAB9 | - | - | C | - | - | - | - | G | - | - | - | - |
| MAS6 | - | - | C | - | - | - | - | G | - | - | - | - |
| MAS10 | - | - | C | - | - | - | - | G | - | - | - | - |

MAB = *M. abscessus*; MAS = *M. massiliense*. The base substitution site is underlined for each codon sequence. The hyphen indicates that the base is the same in the type strain sequence *Mycobacterium* *abscessus* ATCC 19977. The type strain *M. abscessus* is intrinsically susceptible but has inducible resistance to macrolides. T28 is associated with inducible-macrolide resistance. Bases of DNA: Adenine (A), Guanine (G), Cytosine (C), and Thymine (T).

Among the six macrolide-resistant isolates, MAB4 showed the most number of nucleotide substitutions in the *erm*(41) gene. MAB4 is also the only isolate that showed inducible resistance to both azithromycin and clarithromycin earliest on the 7^th^ day. Such polymorphisms include nucleotide substitutions in positions A238G, C253A, A255T, and A312C, all of which lead to missense mutations. Additionally, all isolates harbored the T28 sequevar which is associated with inducible resistance to macrolides. Aside from harboring a thymine at position 28 (T28), the remaining isolates, except for MAB4 and MAB7, showed base substitution at position 168 (G168C). Nevertheless, this only resulted in silent mutation. In addition, base substitution at position 255 (A255G) was found in all isolates, except for MAB4, which also resulted to a silent mutation, thus no observable effect on the organism’s phenotype.

The 836 bp amplicons of the *rrl* gene of macrolide resistant isolates of *M. massiliense* did not show mutations in positions 2057, 2058 and 2059 which are commonly associated with acquired resistance (Wallace *et al*., 1996; Bastian *et al*., 2011; Maurer *et al.*, 2014). Hence, the macrolide resistance shown in these isolates in the study can be attributed to its T28 sequevar in the *erm*(41) gene. There were three *M. abscessus* isolates that showed inducible resistance to macrolides which is attributed to the T28 sequevar of the *erm*(41) gene.

Sequence analysis of the *gyr*A quinolone resistance-determining region (QRDR) revealed only one nucleotide substitution(T252C) present in *M. abscessus* complex isolates, while four nucleotide substitutions (T1443C, C1458T, C1479A, C1494T) were observed in the *gyr*B QRDR (Table 5). Nevertheless, all observed nucleotide substitutions in the *gyr*A and *gyr*B QRDR only caused silent mutations, which do not cause observable effect on the organism’s phenotype. However, it was also observed that all fluoroquinolone resistant isolates of *M. abscessus* complex showed the presence of an alanine residue at position 83 (Ala-83) in the *gyr*A QRDR. The presence of Ala-83 is different from those reported in susceptible bacteria harboring a serine residue at position 83 (Ser-83) (Yoshida *et al.*, 1990). Moreover, all ciprofloxacin resistant isolates harbored an arginine residue at position 447 (Arg-447), and an asparagine residue at position 464 (Asn-464) in the *gyr*B QRDR. These amino acid substitutions are different from those found in susceptible bacteria harboring a lysine at position 447 (Lys-447) and a serine at position 464 (Ser-464_ (Yoshida *et al.*, 1991). These results are in accordance with other studies (Esfahani *et al.*, 2016; de Moura *et al.*, 2012; Guillemin *et al.*, 1998; Guillemin *et al.*, 1995;), showing that substitutions of amino acids at these positions are associated with acquired resistance to fluoroquinolones. It is hypothesized that the presence of these amino acids could decrease the interaction between the quinolone drug and the gyrase A and B subunit-DNA complex (Guillemin *et al.*, 1998).

**Table 5.** Single nucleotide polymorphisms (SNPs) observed in the quinolone resistance-determining region *of gyr*A and *gyr*B of fluoroquinolone-resistant isolates of *M. abscessus* and *M. massiliense*

| Study ID | *gyrA* gene | *gyrB* gene |  |  |  |
| --- | --- | --- | --- | --- | --- |
|  | T252<br><u>TAT</u><br>(Tyr) | T1443<br><u>GGT</u><br>(Gly) | T1458<br><u>TTC</u><br>(Phe) | C1479<br><u>CGC</u><br>(Arg) | C1494<br><u>AAC</u><br>(Asn) |
| MAB1 | C | C | T | - | - |
| MAB5 | C | C | T | A | - |
| MAB6 | C | C | T | - | - |
| MAB8 | C | C | T | - | - |
| MAB9 | - | - | - | - | T |
| MAS6 | - | - | - | - | T |
| MAB10 | C | C | T | - | - |
| MAS8 | C | C | T | - | - |

MAB = *M. abscessus*; MAS = *M. massiliense*. The base substitution site is underlined for each codon sequence. The hyphen indicates that the base is the same in the type strain sequence *Mycobacterium abscessus* ATCC 19977. Bases of DNA: Adenine (A), Guanine (G), Cytosine (C), and Thymine (T).

To our knowledge, this is the first study to assess the drug resistance profile including determination of the genetic mechanisms of drug resistance of *M. abscessus* complex in the Philippines. Therefore, the findings of this study presented a potential area of interest about nontuberculous mycobacteria (NTM) that need to be explored.

Based on the findings of the study, correct identification of *M. abscessus* complex subspecies is important because their responses to anti-TB medications vary from one subspecies to another. Isolation and identification of NTM species, which are not parts of the routine procedure in most clinical microbiology laboratory in the Philippines is important to understand the issues on NTM in clinical samples.

## ACKNOWLEDGMENTS

This research was supported by a grant from the Department of Science and Technology through the Accelerated Science and Technology Human Resource Development Program – National Science Consortium (ASTHRDP-NSC).

